# A systematic scoping review of the ethics of contributor role ontologies and taxonomies

**DOI:** 10.1101/2022.08.29.505654

**Authors:** Mohammad Hosseini, Bert Gordijn, Q. Eileen Wafford, Kristi L. Holmes

**Affiliations:** Department of Preventive Medicine, Northwestern University Feinberg School of Medicine, Chicago, Illinois, USA; Institute of Ethics, Dublin City University, Dublin, Ireland; Galter Health Sciences Library and Learning Center, Northwestern University Feinberg School of Medicine, Chicago, Illinois, USA

**Keywords:** Contributor Roles, Contributor Role Ontology and Taxonomy, Ethics, Authorship, Research Integrity

## Abstract

Contributor Role Ontologies and Taxonomies (CROTs) provide a standard list of roles to specify individual contributions to publications. Due to the recent uptake of CROTs – the CRediT taxonomy in particular– researchers from different disciplinary backgrounds have anticipated a positive impact on ethical issues related to the attribution of credit and responsibilities. Yet, they have also voiced concerns about CROTs shortcomings and ways in which they could be misunderstood or misused and have provided suggestions to improve them. These discussions have never been collated and consolidated. To fill this gap, the current scoping review collates and explores published viewpoints about the ethics of CROTs. Ovid Medline, Scopus, Web of Science, and Google Scholar were searched. In total, 30 papers met the inclusion criteria and were subsequently analyzed using an inductive approach. We identified eight themes and 20 specific issues related to the ethics of CROTs and provided four recommendations for CROT developers: 1) Compile comprehensive instructions that explain how CROTs should be used and that note common pitfalls of employing them in practice; 2) Improve the coherence of used terms, 3) Provide translations of roles in languages other than English, and 4) Communicate a clear vision about future development plans.

## Introduction

Contributor Role Ontologies and Taxonomies (CROTs) are recent innovations developed to address ethical issues associated with the attribution of credit and responsibilities in scholarly publications. By providing a standard list of roles to specify individual contributions to publications, CROTs aim to enhance transparency and consistency about reporting diverse research tasks and contributions to the work leading to a publication, thereby improving the attribution of credit and responsibilities(Brand et al. 2015). The uptake of CROTs has not been limited to journals. CROTs have also been adopted by repositories (e.g., DARIAH-DE, Zenodo) and universities (e.g., University of Glasgow), and are among recommended solutions to improve research assessment, promotion, and funding processes (Hosseini et al., 2022a). Accordingly, CROTs are becoming a pivotal part of scholarly publications and associated workflows, as they are beneficial not only to researchers (e.g., to receive the deserved credit for their contributions) but also for publishers and their editorial teams (e.g., to enhance understanding about who did what in relation to a publication and who is responsible for each task), funding organizations (e.g., to clarify who benefitted from provided funds), universities’ administration and research offices (e.g., to improve tenure and assessment processes) and libraries (e.g., to enable indexing publications based on involved contributions) (Hosseini et al., 2022b). Currently, the Contributor Role Taxonomy (CRediT) is widely adopted (by journals) and used (by the academic community), has been formalized as an ANSI/NISO standard (NISO, 2022a; NISO, 2022b); and has also been integrated into ORCID research identifiers, thereby allowing ORCIDs to be directly linked to the roles (Demain et al., 2021).

Thus far, no published review has systematically analyzed CROTs from an ethical perspective (Hosseini & Gordijn, 2020), which given their growing significance in scholarly publications is a needed area of work. Indeed, with the steady increase of collaborative and international research (National Science Board, 2018), and a continuous rise of the number of diverse roles required in research projects, exploring the ethics of CROTs (i.e., the relevant obligations, values, and virtues in relation to CROTs) increases the likelihood of their ethical development and deployment (Hosseini & Gordijn, 2020). Furthermore, by having ethical issues categorized under specific themes, future research will be provided with an indication of most frequently discussed themes in relation to CROTs.

The first iteration of this review was conducted in 2020 as an exploratory search of the literature about CROTs and was presented in M.H.’s PhD thesis, which was supervised by B.G. and concluded in May 2021 at Dublin City University, Ireland (Hosseini, 2021). However, this effort was unsystematic, limited in scope (only Google Scholar and Web of Science were searched), and lacked input and validation of a librarian and a digital infrastructure expert. To address these limitations, a librarian (Q.E.W.) and a digital infrastructure expert (K.L.H) were invited to collaborate in this scoping review, which endeavors to answer the following questions:

- What ethical issues of CROTs are discussed in the literature?
- What ethical themes can be identified in the current debate about CROTs?

## Methods

After developing a comprehensive search strategy, a scoping review protocol was drafted and revised when necessary (Hosseini et al. 2022c). The search strategy incorporated “contribution”, “attribution” and “authorship” along with keywords and controlled vocabulary terms for CROTs. Terms for CROTs included both descriptive terminology and specific CROT models. We adapted the search strategy to Medline (Ovid), Scopus (Elsevier), and Web of Science (Thomson Reuters). No limitations were placed on document type or publication date. We also searched Google Scholar for additional studies. The bibliographic database searches were performed on January 27, 2022 and identified 979 documents (Full search strategies are available in Appendix 1 and the PRISMA-S checklist is available in Appendix 2).

Upon de-duplication, 576 documents qualified for screening. Titles and abstracts were screened, resulting in the removal of 428 documents based on the following exclusion criteria:

- Non-English sources
- The application of ontologies and taxonomies in areas other than specification of contributions in scholarly collaborations
- CROTs intended for non-scholarly collaborations
- Publication types without a full text

Consequently, 148 documents were deemed eligible for full-text reading. After reading the selected full-text articles, 118 documents were excluded because they did not meet all three inclusion criteria:

- Must mention CROTs that are used to specify contributions to scholarly output (e.g., CRediT, TaDiRAH, CRO, DataCite contributorType).
- Must mention the application of CROTs in reporting contributions made to different scholarly outputs (e.g., manuscripts, research protocols, software).
- Ethical issues (i.e., obligations, values, or virtues) in relation to CROTs should be discussed to a significant extent as determined by an ethics expert (M.H.).

Subsequently, 30 documents that met all three inclusion criteria were deemed eligible for content analysis (Figure 1).

**Fig 1.**
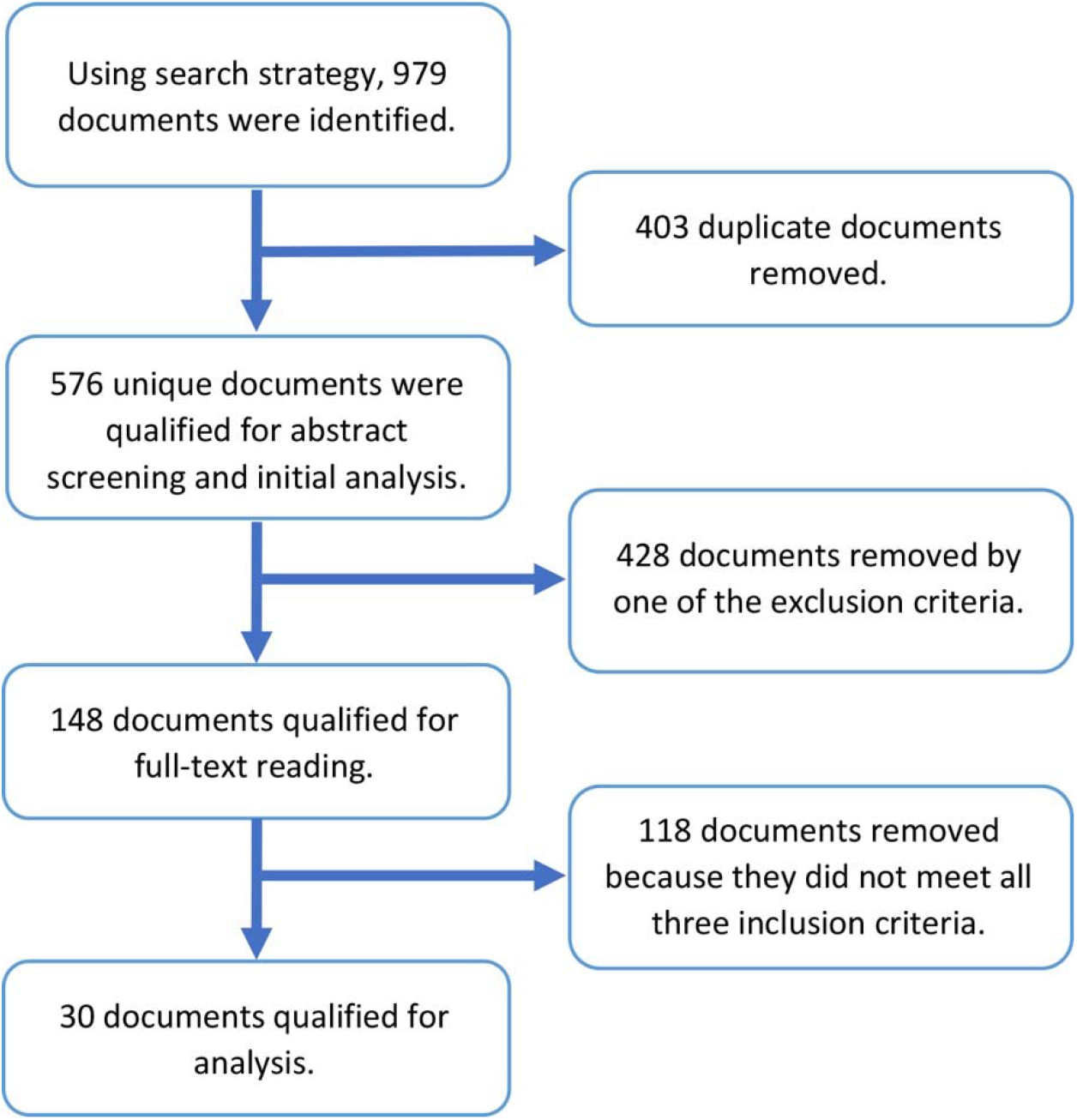
The selection process deemed 30 documents eligible for content analysis.

To analyze the sample, five randomly selected (via computer-generated numbers) documents were examined using an inductive approach, which entails labelling sections that describe an ethical issue followed by subsuming labels under ethical themes, and finally, tagging respective documents with relevant themes (Thomas, 2006). Choosing this method was motivated by the fact that documents that discussed CROTs, reflected views and perspectives of their authors about how CROTs should be used and were not always exploring the ethics of CROTs in a systematic or structured manner. After reading the first five documents, thirteen initial labels were created [M.H.] and improved by co-authors (B.G. and K.H.). Using these labels, the rest of the sample (consisting of 25 documents) was read and labelled, and where necessary, new labels were created. After charting the data in Microsoft Excel, a draft report was developed [M.H.] and shared with co-authors for debate (B.G. and K.H.). This resulted in creating eleven new labels, merging three labels and reducing overlap, bringing the total number of labels to 20. These labels were then subsumed under eight ethical themes. Upon rereading the entire sample, the titles of some issues were changed, but no new themes or issues were created. Eventually, all documents were tagged with one or more themes (Figure 2).

**Fig 2.**
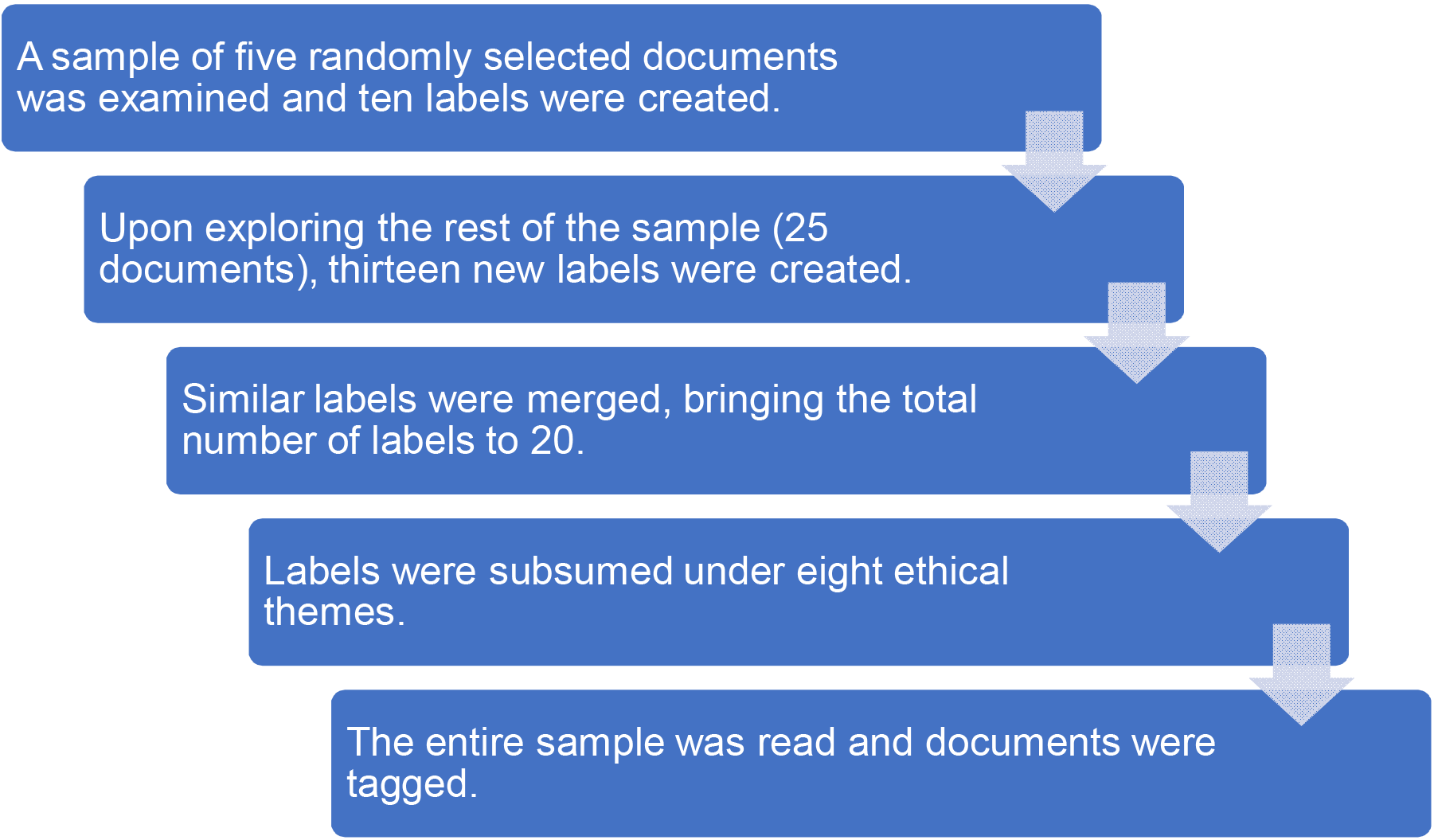
Summary of the tagging process.

## Results

### Characteristics of the sample

#### Document types

Of the 30 documents eligible for full text analysis, 22 were peer-reviewed research articles and the rest were commentary, book chapters and conference papers (Figure 3).

**Fig 3.**
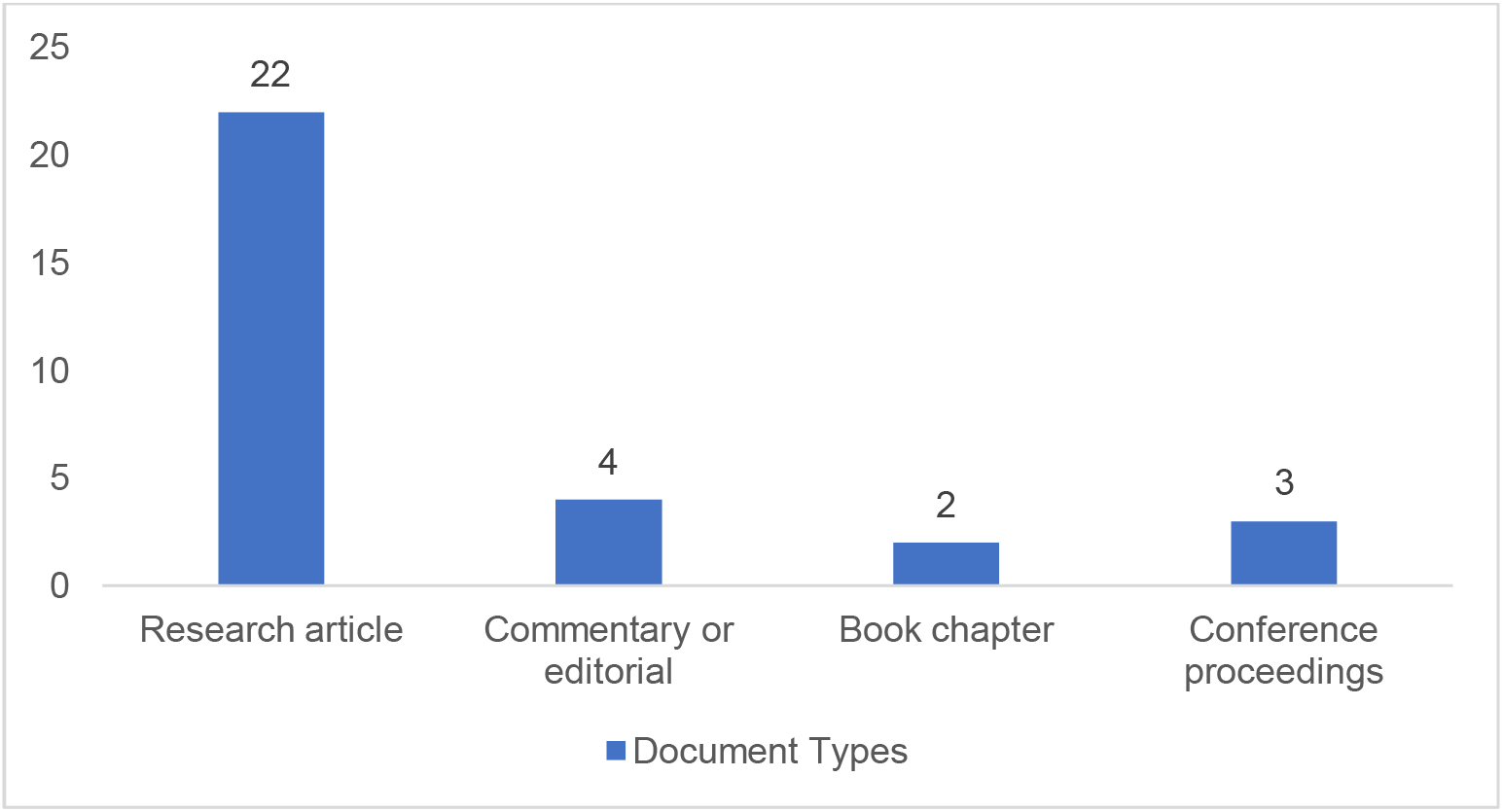
Document types of the eligible items

#### Publication trend over time

The 30 selected articles had been published between 2014 and 2021, and included 10 published in 2021 (Figure 4).

**Figure 4.**
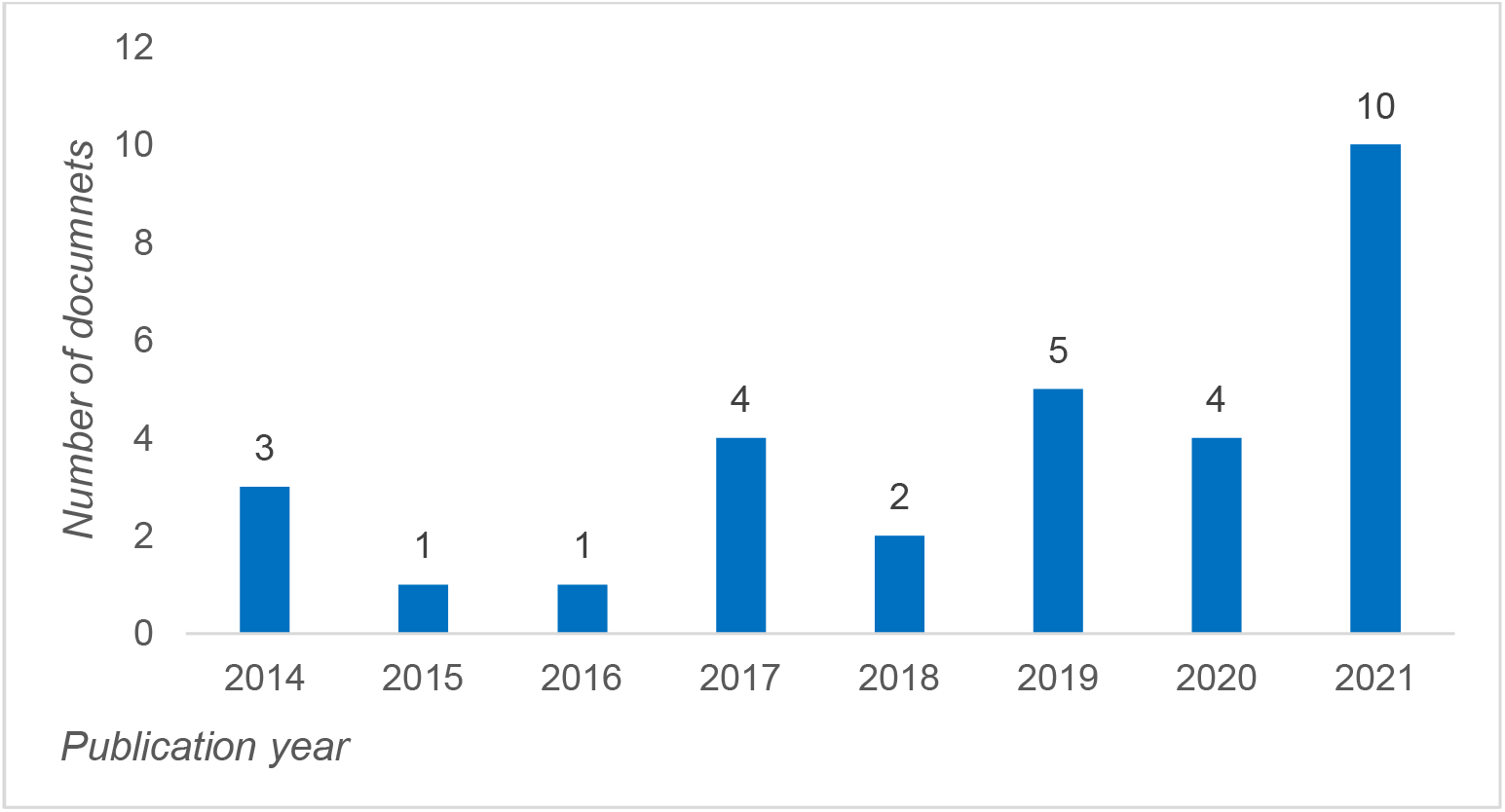
Publication year of the eligible items.

#### Most common ethical issues and the associated themes

Analyzing the sample resulted in identifying eight ethical themes (Figure 5), which encompassed 20 specific ethical issues. In what follows, ethical issues related to each theme are described.

**Fig 5.**
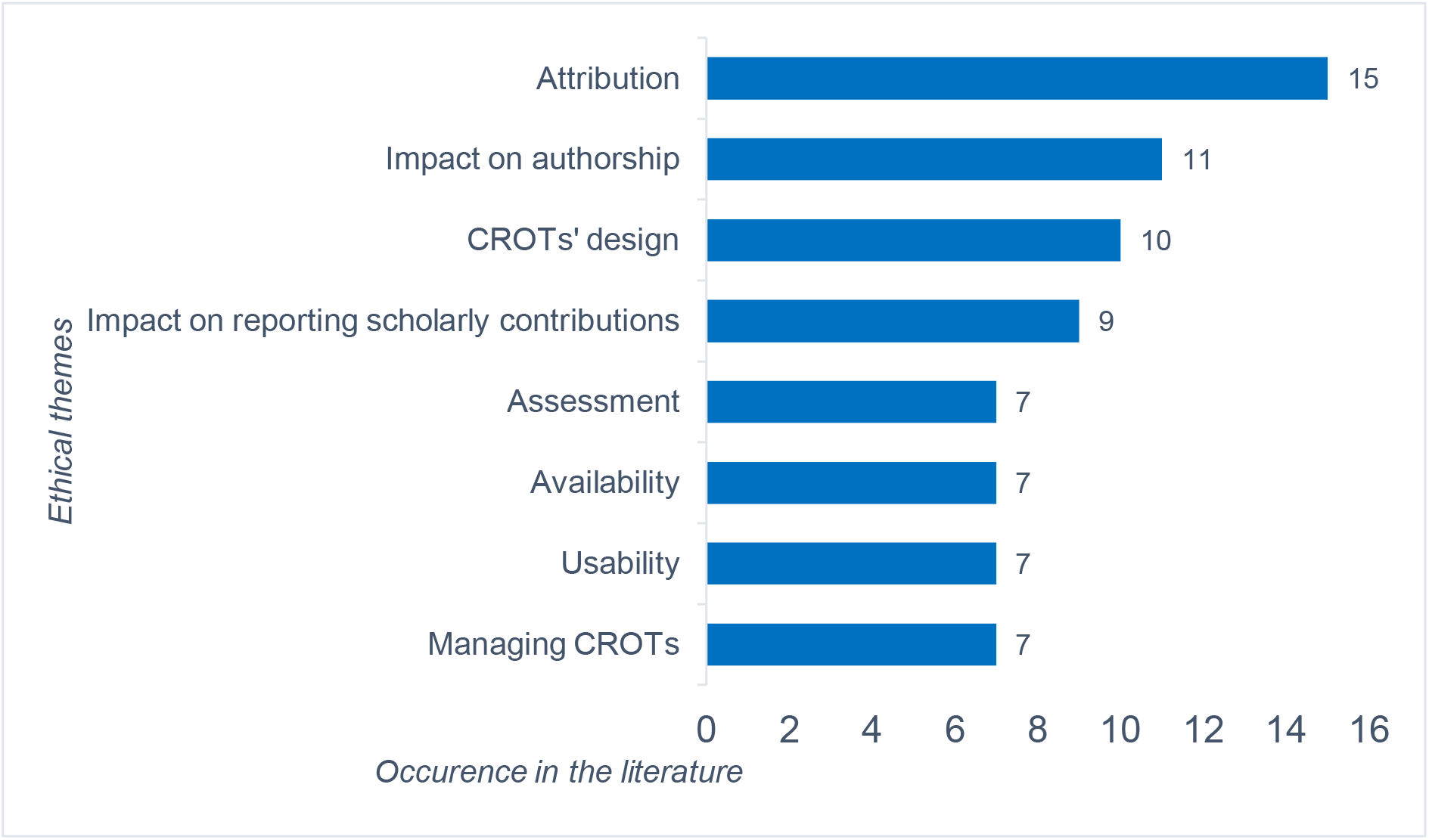
Ethical themes ranked based on their occurrence in the literature. Some documents are tagged with more than one theme.

### 1. Attribution

Attribution issues are about improving CROTs’ lists of roles and definitions, extending output types that should be captured by CROTs and using CROTs in specific contexts, all of which directly impact the scope of credit attributions and the extent to which specific disciplinary nuances are captured by CROTs.

#### Issue 1) Improving CROTs’ lists of roles and definitions

CROTs’ list of roles should evolve “as science and the types of contributions that may become less or more important change” (Allen et al., 2019, p. 73). Indeed, keeping lists current to meet evolving user needs is paramount and among factors that promote the use of CROTs (Vasilevsky et al., 2020). Although Ding et al. (2021, p. 7578) and colleagues asserted that CRediT roles “cover almost the whole process of scientific research of most disciplines”, several publications mentioned specific roles that should be added to CRediT, perhaps because as its developers note, “CRediT was initially developed and tested as a taxonomy for life and physical sciences” and thus, may not fit all other disciplines (Allen et al., 2019, p. 73). For instance, Alpi and Akers (2021, p. 362) noted that “CRediT does not currently describe all roles that librarians play”. Steele et al. (2021) examined the ability of CRediT to capture the contribution of medical writers to reports of Randomized Controlled Trials (RCTs) in dermatology. They found that in only 1% of published RCTs a distinction is made between copyediting and preparing the first draft (Ibid.).

While Dappert et al. (2017)made a general suggestion to define roles more clearly, others made more specific suggestions. For instance, Matarese and Shashok (2019, p. 6) claimed that CRediT “ignores” some non-author contributions including technical support, translation and editing the manuscript. They provide specific suggestions to improve three of CRediT’s current roles (Investigation; Writing—Original Draft; Writing— Review & Editing) and add two new roles (Technical support; Translating or editing the manuscript, as non-author). Zhang et al. (2019, p. 6) noted that some important roles in RCTs such as “randomization, patient enrollment, and follow-up are not clearly defined” by CRediT, and suggested Contributor Roles Taxonomy for Randomized Controlled Trials (CRediT-RCT) that includes ten roles for conducting an RCT (Conceptualization, Funding Acquisition, Project Administration, Site Principal Investigator, Statistical Analysis Plan, Investigation, Data Curation, Formal Analysis, Writing—Original Draft, Writing—Review & Editing). The role of software development for research and CRediT’s shortcomings in capturing involved nuances is highlighted by Alliez and colleagues (2020). They argued that since software development in research involves “significant innovations”, qualified human intervention and qualitative information are necessary to ensure accurate reporting of contributions (Ibid., p. 48). Accordingly, they suggested nine finer categories for better representation of involved tasks (Design, Debugging, Maintenance, Coding, Architecture, Documentation, Testing, Support, Management).

#### Issue 2) Extending output types

Dappert et al. (2017) noted that CROTs’ scope should be extended beyond peer-reviewed journal articles. While supporting this suggestion, Vasilevsky et al. (2021, p. 35) highlighted that since output types such as datasets, software, and research protocols “have less well-established workflows to collect and present structured metadata”, this is not easily achieved. Similarly, although two papers specifically noted that capturing contributions to datasets should be improved (McNutt et al., 2018; Mongeon et al., 2017), others highlighted the lack of a robust link between datasets and all of their contributors as a factor that “inhibits re-use, verification of research, and detection of scientific fraud” in datasets (Fenner et al., 2015, p. 309). The absence of robust links between contributors and datasets could be explained by the wide range of contributors to datasets (e.g., governments, nonresearch organization, companies, and other professionals), which often include groups and teams as authors *in addition* to individual authors. Since there is no ORCID equivalent for groups and teams, such practices reduce consistency and machine-readability of attributions (Mongeon et al., 2017; Dudek et al., 2019). Although adding groups and teams to the authorship list is also common in peer-reviewed papers, none of the explored items in the sample mentioned this issue as a challenge.

#### Issue 3) The use of CROTs in specific contexts

Extending the application of CROTs to other object types could be particularly challenging in some contexts. According to Bliss et al. (2020, p. 2), in creating digital outputs such as online dictionaries, oral story databases or linguistic atlases, some contributors might only be “involved for a very short time, as short even as just one recording session or consultation”, but others’ contributions might extend to years. This dynamic justifies devising *macro-credits* and *micro-credits* to account for discrepancies in terms of the time contributors were engaged in a project. Bliss and colleagues further noted that in their evolving project, which constantly attracts new contributors and contribution types, macro and micro credits are needed. While macro-credits are high-level “general contributor roles such as editor, director, web design, data processing, and funding”, micro-credits are attached to individual items such as dictionary entry and audio files, and “include more specific contributor roles such as speaker, storyteller” (Ibid., p. 13).

Alliez et al. (2020, p. 47) highlighted the complexities of capturing contributions made to software and question the accuracy of taxonomies that flatten out all involved contributions “on the simple role of software developer”. McLaren and Dent (2021, p. 53) noted that some outputs such as “design protypes, experimental data sets, or methodologies designed to test a hypothesis” are not captured in a digital format during research workflows and sometimes emerge from intangible and immeasurable contributions (e.g., experience), thereby making them complicated to capture by CROTs.

### 2. Impact on authorship

Since CROTs are currently used in parallel with authorship bylines, they are discussed within debates about ethics of authorship and sometimes promoted as a solution to resolve these ethical issues.

#### Issue 4) Authorship disputes

Some publications suggest that using CROTs (and CRediT in particular) minimizes the likelihood of specific ethical issues of authorship. Using CRediT is believed to not only reduce authorship disputes (Brand et al., 2015), but also “support research institutions and authors to *resolve* author disputes by providing more transparency around individual author roles and responsibility” (emphasis added) (Allen et al., 2019, p. 72). Holcombe (2019) anticipates that since CRediT is implemented with checkboxes to choose contributions (as opposed to authorship criteria that requires reading large amounts of text and complicated inferences), it is likely to induce better engagement from researchers because navigating checkboxes is easier than reading authorship criteria. However, Matarese and Shashok (2019) noted that checkboxes induce authors to choose more roles than they would declare in a free-text form.

#### Issue 5) Honorary authorship

Teixeira da Silva (2021) and Holcomb (2019) suggested that using CRediT reduces honorary authorship because it reduces ambiguity about contribution types. However, this view is challenged by Larivière et al. (2021, p. 124)who argued that researchers could also adopt practices like ghost, guest, and gift authorship when using CROTs. They further concluded that CROTs “systematic description of work does not, therefore, preclude invisibility, but only displaces it elsewhere. Consequently, it leaves ghostwriting of articles and potential honorary contributorship in the backrooms of scientific research” (Ibid.).

#### Issue 6) Authorship order

Using CRediT is believed to reduce ethical challenges regarding authorship order as well. Without alluding to *how* using CROTs will influence author order across disciplines (where order is strongly guided by specific norms), McNutt et al. (2018, p. 2559) claimed that adopting CRediT would “help alleviate some of the confusion across disciplines and cultures regarding the meaning of author order”. Using CRediT is also considered as a reasonable strategy to “minimize interpretation of [authorship] order and variability in perceptions of roles” (Poirier et al., 2021, p. 226).

#### Issue 7) Authorship standards

Although McNutt et al. (2018) and Holcombe, (2019)explicitly noted that CRediT improves current authorship standards from an ethical perspective, some explained why this claim might not be always true. For example, Bliss et al. (2020) argued that CRediT does not help distinguishing those who should be listed in the author byline from those who should be acknowledged, and, Matarese and Shashok (2019, p. 6) it is “only applicable to byline authors”. This could, in part, be because CRediT (as a model) does not resolve questions “about the quantity and quality of contribution that qualify for authorship” (Larivière et al., 2021, p. 124), and while some of its roles overlap with elements of authorship, they “do not fully encapsulate in any single role, the concept of authorship” (McLaren & Dent, 2021, p. 53). Furthermore, CRediT does not provide clarity on one of the most challenging aspects of defining authorship, namely the notion of “substantial contribution” (Smith & Masters, 2017, p. 253).

### 3. CROTs’ design

Ethical issues about CROTs’ design pertain to the overall architecture and design of CROTs that ultimately affect an ethical attribution of credit and responsibilities.

#### Issue 8) Open or controlled vocabularies

Open-ended lists allow the addition of new roles and thus, essentially accommodate the recognition of any task deemed relevant in a project, but controlled lists only entail a certain number of tasks and unless updated, only allow recognizing these tasks. In their introduction of the CRediT taxonomy, Brand et al (2015, p. 154) stressed that “a controlled vocabulary of contributor roles” is needed within the scholarly ecosystem. Craig’s (2018, p. 46) citation of suggestions provided by the UK Academy of Medical Sciences explained the rationale for this need: “So that individuals encounter the same system in all work, whether as a researcher or an appraiser”. Those in favor of open vocabularies, however, advocated for “fast changing” fields in terms of used techniques and objects, e.g., digital humanities (Dombrowsky & Perkins, 2014, para. 2) or projects that involve a “continuously evolving” set of roles both at the project and item level, as well as the preference to prioritize “the wishes of the contributors of each project over perceived consistency” (Bliss et al., 2020, p. 13).

Although both Larivière et al. (2021) and Borek et al. (2021, p. 325) noted that a CROT’s list of roles can *never* be complete, the latter added that they do not have to be complete but instead should be “open for extensions without the need to revise existing definitions”. The open-endedness of list of vocabularies is also implied in design requirements suggested by Sauermann and Haeussler (2017, p. 9) who stressed that CROTs should be flexible to accommodate nuances across projects and fields and, also, “anticipate changes to scientific activity, such as growing team size and specialization, automation and commoditization of certain research activities, as well as broader participation by nonprofessional scientists”.

#### Issue 9) Strategies for selecting terms for a vocabulary

Strategies used for choosing terms for a vocabulary, affect how vocabularies are understood and used in practice, thereby affecting the attribution of credit and responsibilities. While specific terms support precision, broad terms support “consistent application, collocation and recall” (Borek et al., 2014, p. 182). Furthermore, selected terms should be reconciled with existing taxonomies (e.g., tags currently used by repositories and libraries) and strike a balance between theoretical correctness and commonly used terms “(visualization + geospatial coordinates object vs. mapping)” (Ibid.). Selecting terms for vocabularies also affects the evolution of CROTs, because sometimes decisions must be made about whether a role is “distinct enough to stand on its own as a separate category (e.g. storage vs. storage and dissemination)” or it can be subsumed under existing categories (Borek et al., 2016, p. 5).

Selecting terms for roles that are inherently related and/or roles that are conducted by the same person are discussed as well. For instance, Zhang et al. (2019, p. 6)suggested that in RCTs, roles such as formal analysis and software (both among CRediT roles) cannot always be described with unique terms because “statisticians need to use software to perform formal analysis”. Challenges of updating terms in CROTs are briefly discussed in relation to TaDiRAH, whose developers suggested using “the property skos:closeMatch” to create a mapping between old and new terms (Borek et al. 2021, p. 328).

### 4. Impact on reporting scholarly contributions

These ethical issues are concerned with the impact of using CROTs on reported contributions to scholarly work.

#### Issue 10) Strategic and opportunistic use of CROTs

Larivière et al. (2021, p. 124)noted that since contributions and gained credit are inevitably tied to the reward system of science and the *symbolic capital* in academia, goal displacement (i.e., behavioral modification to “meet certain requirements, norms or incentives”) might happen upon wider adoption of CROTs (2021, p.). As an example, they suggested that the disproportionately high number of *PLOS Medicine* authors assigned with CRediT’s roles of “Writing—Original Draft” and “Writing—Review & Editing” may be explained by authors’ attempt to demonstrate adherence to the ICMJE authorship criteria (as the ICMJE’s second criterion explicitly requires a contribution to the writing process). Furthermore, they noted the risk of financial conflict of interests being disguised when CROTs are in publications that report industry-academia collaborations. From an industry-partner perspective, authorship implies *intellectual intervention* whereas contribution to *tasks* might help “avoid allegations of conflicts of interest” (Ibid.).

Three publications mentioned the impact of CROTs on inaccurate reporting of contributions. Matarese and Shashok (2019, p.8) noted that since CRediT is only used to describe authors’ contributions (thereby omitting non-author contributors), it could induce authors to “unintentionally yet misleadingly shift credit for certain tasks from the people who did the work to people named in the byline as authors”. Larivière et al. (2021, p. 124) expressed concerns about disciplinary differences in terms of interpreting contributions, and the *presumed* link between the actual labor and its indicators, speculating that the discrepancy between the two could be further obscured by researchers’ strategic behavior, e.g., favoring “good working relationships”. Sauermann and Haeussler (2017) highlighted that using CROTs may encourage researchers to crowd into tasks that are considered more valuable and dissociate themselves with tasks considered less important or those with a greater potential for risk of error.

#### Issue 11) Responsibility and accountability

A survey to investigate contributions made by undergraduate assistants to research projects suggested that since collaborators with different levels of seniority might have dissimilar assumptions about the “level” as well as “range of responsibilities over time”, mentors and students may sometimes have different impressions of individual responsibilities (Honoré et al., 2020, p. 42). For example, when conducting *Data curation* and *Investigation* tasks (often involving “repetitive work guided by structured procedures and protocols”), both mentors and student researchers assume high levels of responsibility for students. However, when it comes to tasks such as *Validation* or *Writing*, a discrepancy is reported between assumed responsibilities (Ibid.).

Two publications mentioned CRediT and the ICMJE criteria in the context of responsibilities. McLaren and Dent (2021, p. 53) noted that while authorship credit “carries additional layers of rights, responsibilities, and ultimate accountability for the work” and is also subject to stringent tests such as the ICMJE criteria, CRediT is only concerned with acknowledging (some of the) contributions. Teixeira da Silva (2021, p. 1118) noted that CRediT is an improvement to the ICMJE recommendations “increasing the transparency of author contributions and stating with greater clarity the contribution of each author, potentially fortifying accountability”.

#### Issue 12) Significance and extent of contributions

Two publications highlighted CRediT’s inability to rate different tasks based on their significance for a project, which results in ambiguities about the overall significance of individual contributions. While Ding et al. (2021) noted that knowing each role’s significance in relation to the whole work helps differentiate their importance, the argument provided by Smith and Masters (2017) — i.e., that the importance or value of roles varies per project and context, implies that this clarification cannot be a default assumption built into CROTs (2017). Furthermore, Simth and Masters (2017, p. 253) provided an analogy to the notion of substantial contribution in using authorship definitions and suggest that when using CRediT, “it remains unclear how much an individual must contribute” to qualify as a contributor. They noted that since being *fully inclusive* results in lengthy lists of names “including those individuals who provide minor contributions”, some form of quantification “is, and will always be, necessary” (Ibid.). Nevertheless, Alliez et al. (2020) claimed that attempts to quantify contributions, even if automated (e.g., to capture contributions to software development), could be gamed to inflate contributions to an individual’s benefit.

### 5. Assessment

The attributed credit for conducting research-related tasks is ultimately reflected in resumes and used for assessing researchers in academic hiring and promotion processes. These ethical issues are concerned with the implications of using CROTs in academic assessments.

#### Issue 13) Improving academic assessment and evaluation

While Smith and Masters (2017) noted that CROTs are unlikely to be used as metrics of performance without authorship, others argued that they may facilitate “the development of contribution-based indices to complement authorship-based indices” (Sauermann & Haeussler, 2017, p. 9) or allow developing altogether “new metrics of individual research productivity and impact” (Alpi & Akers, 2021, p. 362). Whether as standalone or in combination with authorship, these changes require the development of infrastructure to integrate CROTs into funding agencies’ submissions and progress reports and institutional hiring and assessment workflows (Vasilevsky et al., 2021). Another aspect where CROTs can benefit research assessment pertains to the peer review process. Integrating CROTs into review processes could improve efficiency by means of connecting reviewers to specific contributions based on proficiency and expertise (Ilik et al., 2018; McNutt et al. 2018; McLaren & Dent, 2021).

#### Issue 14) Improving equity and diversity in academic work and evaluation

Two publications claimed that since using CROTs improves Meta Science (i.e., the science of science), their wide adoption could provide new insights about the distribution of labor in academia and help addressing gender and diversity challenges (Alpi & Akers, 2021; Allen et al. 2019). In fact, Larivière et al. (2019) use of the CRediT roles of articles published in PLOS journals between 2017 and 2018 is an example of how CROTs could be used in Meta Science (these authors note that the quality of such studies is highly dependent on the accuracy of reported contributions). Among other conclusions, Larivière et al. (2021, p. 119) noted that “men are more likely to conduct tasks associated with seniority, such as funding acquisition and supervision (30% more likely than women), contributing resources, software, conceptualization, and project administration”.

### 6. Availability

CROTs’ availability pertains to their accessibility for different members of the academic community and discoverability for reuse, both of which affect how credit is distributed among different cohorts.

#### Issue 15) Accessibility

Since inaccessible or partially accessible CROTs might negatively impact different user groups such as researchers and institutions, conformity of CROTs and contributors’ data to FAIR principles (Findable, Accessible, Interoperable and Reusable) is essential. This could be demonstrated, for example, by using unique resource identifiers “to uniquely identify the concepts”, “using W3C Standard SKOS and the usage of CC0 1.0 licence” (Borek et al., 2021, p. 330). Furthermore, different forms of accessibility were mentioned in two papers. Although Craig (2018) emphasized the electronic accessibility of contributor information (e.g., in journals’ websites), Das and Das noted that such developments cannot be implemented “in the print copy” of journals (2020, p. 29).

Availability of CROTs has also been discussed in terms of translating roles and their definitions into languages other than English. For instance, TaDiRAH’s translation into German, French, Spanish, Portuguese, Serbian, Italian and Norwegian will not only “ensure technical and professional mentoring for the use and reuse” (Borek et al., 2021, p. 330), it will also make it more accessible to more users across the globe (Borek et al. 2016). Indeed, increasing the visibility of contribution statements requires reducing access barriers and processing costs, and could be further facilitated by editors, funding agencies, administrators, and database providers (Sauermann & Haeussler, 2017).

#### Issue 16) Machine-readability

Machine-readable contributions are readily discoverable and reusable (Vasilevsky et al., 2021; Craig, 2018)], improve availability of CROTs to more users and extend the application of CROTs to a “variety of contexts” (Borek et al., 2016, para. 38). This is further discussed by McNutt et al. (2018, p. 2559) who noted that the availability of contributor information in both machine- and human-readable forms facilitates the “transparency of author contributions in different contexts, via syndication, indexing, and abstracting services, and possibly future applications across journals”. When contributor information is available in a machine-readable and consistent format, “contributions will transition from hearsay to quantifiable evidence” (Ibid.) [assuming, of course, that the data are accurate].

### 7. Usability

Ethical issues about usability include suggestions that affect how CROTs are used by the academic community, thereby impacting the attribution/presentation of attributed credit and responsibilities.

#### Issue 17) Improving usability

Some suggestions to improve usability require technical development, such as linking CRediT roles to ORCID records (Craig, 2018; Vasilevsky et al., 2021) and creating free web-based tools that allow journals to keep additional information in “acknowledgments, footnotes, or by other means” (McNutt et al., 2018, p. 2560). McLaren and Dent (2021, p. 53) noted that CRediT is not consistently understood and applied across the board and highlighted supplementary approaches that could complement the CRediT model, e.g., the “Technician Commitment” initiative in the UK. Another suggestion to improve usability is to develop and promote guidelines that describe how contributions should be allocated (Craig, 2018). For example, Larivière et al. (2021, p. 124) suggested that since only 55% of explored articles in a sample of 30,054 included the role of validation, “journals could require validation as a mandatory contribution type for empirical work”.

#### Issue 18) Assigning roles

Three papers discussed role assignment. Brand et al. (2015) and colleagues recommended *corresponding authors* to assign CRediT roles and to provide the review and confirmation opportunity to other contributors. However, in describing how to use CRO, Ilik et al. (2018, p. 7) proposed a different workflow for role assignment, calling it the “claim process”. In this process, which is part of the paper production stage, authors, non-authors, editors and others report their own contributions using the list of roles (instead of the corresponding author assigning these and having contributors confirm). Once contributions are reported, all contributors would confirm reported roles. Another approach is adopted by the Algonquian Language Digital Resources Credit System wherein an additional free-text box allows contributors to describe roles *in their own words* to ensure more transparency (Bliss et al. 2020).

### 8. Managing CROTs

Since contribution types in various research fields change over time, CROTs should evolve and reflect new roles to ensure an ethical attribution of credit and responsibilities across the board. This requires a CROT’s administrators and management board to engage with the academic community.

#### Issue 19) Transparency

The transparency of involved process in the development and revision of CROTs is discussed in two publications. The CRediT developers described the 2012 collaborative workshop held at Harvard University as an event that engaged experts and as a defining moment in the development of the taxonomy (Brand et al., 2015). In contrast, Matarese and Shashok (2019, p. 5) questioned the openness and transparency of this process, noting that this was “an invitation-only meeting”, and that data from the subsequent survey of life science researchers who provided “positive feedback” about the initially suggested taxa were never made available. They also reflected on CRediT’s ownership structure and claim that Elsevier is “the de facto owner of CRediT” (Ibid.). (In further inquiry about this claim, we reached out to the CRediT standing committee and asked for clarification. They disagreed with the claim and replied: “Aries and Elsevier own and sell licensed access to the software that is capable of collecting, storing, and exporting the roles. The role metadata that the system collects are property of the journal” (Personal Communication, 20 April 2022).)

#### Issue 20) Engagement

Two papers introduced open community-developed resources and mechanisms to collect user feedback as useful tools for engagement with the academic community, which also promote collaborative approaches in managing activities related to CRO and CRediT (Ilik et al., 2018; Vasilevsky et al., 2021). Borek et al. (2016) highlighted using similar strategies for maintenance of TaDiRAH and note that versioning and issue tracking features of GitHub have been conducive to their efforts. However, they also highlighted that one challenge of open engagement pertains to receiving contradictory feedback from the community. Borek et al. (2021, p. 330) also introduced the “TaDiRAH board”, which will be responsible for managing activities related to the development of TaDiRAH: “The board consists of the original core team, new developers and other contributors”. Furthermore, this group openly shared their strategy for expanding TaDiRAH to “revise and harmonize the structure and semantics of the model, as well as the concept terms and definitions” based on how it is used by the community (Ibid.).

## Discussion

### Confusion around CRediT’s application and impact

While CRediT is the most widely used among the CROTs (and also discussed more frequently, as shown in this review), its correct application and possible impact on current ethical issues are not always fully clear to those who discuss it. Reviewing the literature showed a discrepancy between what CRediT developers advise and what the community understands in terms of CRediT’s scope and application – some papers in our sample have misread or misunderstood key details about CRediT (e.g., (Alpi & Akers, 2021; Honoré et al. 2020; McNutt et al., 2018)).

CRediT developers have published several journal articles to create awareness about CRediT and its intended use (Allen et al., 2019; Brand et al. 2015) but misinterpretations in the published literature suggest that these instructions have not always been effectively communicated and well-received or perhaps journal articles were misread.

For instance, in an editorial published in *Journal of the Medical Library Association*, Alpi and Akers (2021, p.109) noted: “Together with ICMJE guidelines, CRediT can be used to facilitate conversations and *help determine who merits authorship within collaborative teams* (emphasis added)”. In a different example, Honoré et al. (2020, p. 42) claimed that CRediT was “originally conceived and designed with authorship roles in mind”. However, Brand et al. (2015, p. 154) have refuted both claims when they described the taxonomy’s scope, and specifically highlighted that CRediT’s parallel use with authorship definitions also caused confusion when CRediT was piloted for the first time:

> “Most of the questions that arose concerned confusion over whether the taxonomy was explicitly intended to specify which types of contribution qualify for authorship status, when in fact that was never the intention. As stated in the taxonomy header: The classification includes, but is not limited to, traditional authorship roles. That is, *these roles are not intended to define what constitutes authorship*. Rather, the roles are intended to apply to all those who contribute to research that results in scholarly published works, and it is recommended that all tagged contributors be listed, whether they are formally listed as authors or named in acknowledgements”.

Other examples of misunderstanding pertain to CRediT’s impact on reducing ethical issues related to authorship order. When promoting CRediT, Brand and colleagues briefly mentioned authorship order following a hypothetical situation in a non-existing infrastructure, and stipulated that *assessing* researchers in such a scenario would no longer depend on authorship order:

> “Imagine publishers collecting structured information about contribution in a standard format. Imagine, further, that this information is associated with the article DOI, via CrossRef, and with ORCID author identifiers. We would then have the infrastructure in place to track not only who authored which publications, but also who contributed what to each publication that names the individual as a contributor. With this infrastructure in place, it would eventually be possible to devise more precise, author-centric credit and impact tracking tools, on which the byline order of author names would have no bearing” (Brand et al., 2015, p. 154).

However, McNutt et al. (2018, p. 2559),-cited by 37 and 215 articles as per October 2022 according to PubMed and Google Scholar, respectively, noted: “Adopting the CRediT taxonomy would also help alleviate some of the confusion across disciplines and cultures regarding the meaning of author order”. To back this claim, authors cited Sauermann and Haeussler’s 2017 paper, which did not make such a claim either.

In a different misunderstanding about CRediT’s impact on reducing ethical issues related to authorship order, while citing Brand et al. (2015), Poirer et al. (2021, pp. 225-226) noted: “Adhering to the recommendations from Project Credits [CRediT] by clearly listing contributions of each author may be a reasonable strategy for acknowledging order. This approach would create transparency and minimize interpretation of order and variability in perceptions of roles”.

Examples like these or instances wherein journal instructions suggest using CRediT contrary to what is prescribed by its developers (seeHosseini et al., 2022a for an example about the *PLOS Biology* journal) show that communicating information about CROTs by means of peer-reviewed publications is not always effective. CROTs such as CRediT could be misused and misunderstood in many ways, and researchers will not always have the knowledge necessary to identify, read and understand the instructions embedded in peer-reviewed publications. One way of minimizing these misunderstandings, is to develop reasonably comprehensive and straightforward instructions for users (similar to those developed by the ICMJE).

### Issues relating to CRediT’s current list of terms

Some CRediT roles describe tasks at the project-level while others describe tasks at the paper-level. While one of the ethical issues relate to CROTs’ design pertained to strategies for selecting terms for a vocabulary, none of the explored papers mentioned certain irregularities in CRediT’s current list of terms. Here we discuss two such issues, 1) using both performative and non-performative terms, 2) confounding paper and project-level contributions.

1. While TaDiRAH developers use a coherent and consistent set of terms for activities, arguing that these terms should be “gerundiva [i.e., verbal adjective] to represent the performativity of the activities” (Borek et al. 2021, p. 325), such considerations have not been observed in developing CRediT terms. Consequently, some CRediT terms are non-performative nouns (e.g., resources, software) that strictly speaking, do not represent a specific activity.
2. CRediT currently dovetails roles that describe specific work on a paper (e.g., writing, methodology, validation) with roles that mainly occur at a project level, and that could affect (and be reflected in) several papers (e.g., funding acquisition, project administration). This inconsistency could result in project-level roles being more coveted (and preferred by senior researchers) because they can potentially yield better return on invested time: Suppose project X has eight contributors involved in investigation, data curation and writing, two contributors involved in funding acquisition and one in project administration. If project X results in five papers, those involved in roles such as investigation, data curation and writing would have to make specific contributions in relation to each paper to be listed as a contributor, but those who acquired funding and performed administrative tasks for the *project* would be automatically listed on all papers.

Given CRediT’s quick uptake and a recent study that found that CRediT roles show an unequal distribution of labor and gender differences in terms of conducting specific tasks (Larivière et al., 2021), it is reasonable to expect CRediT developers to improve these irregularities in future iterations.

### CROTs status: complementary or stand-alone?

Are CROTs complementary to existing authorship bylines or would they ultimately become stand-alone solutions? Papers that discuss CROTs are not always clear about whether, and how, CROTs should be employed in tandem with authorship lists. Given 1) legal and 2) ethical challenges of doing away with authorship lists, currently, it seems unlikely (if not impossible) to envision scholarly publications without author lists:

1. Authorship has a legal significance – authors can readily engage in litigation in relation to a publication, whereas non-authors first have to prove that they have a legitimate claim to authorship. Employing CROTs without authorship lists affects “copyrights, intellectual property rights and the process via which these rights are assigned/challenged” (Hosseini, 2021, p. 260).
2. Authors’ names are used for (non-numerical) citations and in reference lists. Without author names, publishers and/or the scientific community must develop and adopt completely new systems of citation and referencing based on contributors’ names. In such a system, ethical issues might occur with regard to the selection of the contributor(s) whose name(s) should appear (at all/first) on intext citations or in the reference lists. Furthermore, while using authorship lists (except those ordered alphabetically), for the most part, it is possible to implicitly infer those who made the highest degree of contribution relative to others (i.e., first author) and those responsible for the overall integrity of the work (i.e., last author). However, CROTs cannot currently address these issues on their own (Ibid., p. 264). Even using the Supervision role cannot always help because projects might have numerous supervisors.

Accordingly, in the foreseeable future, CRediT (and other taxonomies) must be used in parallel with author bylines. Nevertheless, the scope of this parallel use is currently vague and visions to improve or end it are not openly communicated. Specifically, since its developers suggest that CRediT should <u>not</u> be used to define authorship, and, CRediT does <u>not</u> capture the full range of scholarly contributions – even in life and physical sciences for which it was initially intended; it is unclear how the CRediT roles and authorship definitions should be employed beside each other. Answering questions like these would facilitate developing a clearer vision about the function which CRediT and other CROTs could serve in the future of scholarly publications.

### CROTs are not the panacea

When looking at the ethical issues of CROTs, similarities with ethical issues of authorship are noticeable. For example, ethical issues about the attribution of credit and responsibilities when using CROTs are similar to ethical issues about the attribution of authorship credit and associated responsibilities, and ethical issues about CROTs’ design and usability have similar components to ethical issues about the definitions of authorship and their application in different contexts. Given these similarities, one can argue that in discussing scholarly attributions, improving the used model (e.g., authorship definitions, CROTs) and the development of technical workflows or even the creation of a *perfect CROT schema and workflow* may address technical (or syntax) problems but not the human or social component of the problem (arguably much more urgent and challenging to address). Exploring the similarities between ethical issues of authorship and CROTs in future research could lead to new insights about unethical human behavior in attributions as well as challenges of recognizing scholarly work.

This review is limited by the small sample size and does not include debates that took place on the blog-sphere, online forums or in print-only material. Exploring these spaces could be useful for future research. In addition, this review only used and considered English publications about CROTs, and therefore, exploring viewpoints published in other languages could be among possible directions for future research. Conducting empirical research to explore the views of researchers from different disciplinary backgrounds could be another area for future research.

## Conclusions and recommendations

While our review of the debate about CROTs demonstrates a growing interest in CROTs in recent years, the academic community should be mindful about inaccurate published views about them, especially regarding how they should be used and the implications of their adoption. We believe that CROTs should be continuously improved to create better systems of attribution, but overemphasizing CROTs shortcomings could potentially be used to excuse unethical behavior of researchers or perverse institutional incentives, both of which continue to be urgent to address. In what follows we provide four recommendations:

1. **Compile and promote comprehensive instructions that explain how CROTs should be used and that note common pitfalls of employing them in practice.** Given the mentioned ethical issues in terms of CROTs’ impact on authorship and reporting of scholarly contributions, as well as the highlighted inaccuracies in the literature, extensive and specific guidance is needed to orient scholars about the concept of CROTs and guide their responsible use. In particular, communicating the scope and limitations of using CROTs in parallel to authorship bylines seems essential.
2. **Improve the coherence of existing roles**. Our analysis of existing CRediT roles shows that some roles describe tasks at the project-level while others describe tasks at the paper-level. In the long run, these incoherencies might create hierarchies among roles and result in an unfair attribution of credit.
3. **Provide translations of roles in languages other than English**. Currently, only one of the CROTs has roles translated into languages other English (e.g., TaDiRAH). Given the wide international use of CROTs, translation of roles into non-English languages improves availability and inclusiveness of CROTs.
4. **Communicate a clear vision and strategy about future development plans.** Timely communication of future plans could facilitate input and engagement from the community and democratize the process of improving CROTs.

## Appendix 1 Database Search Overview

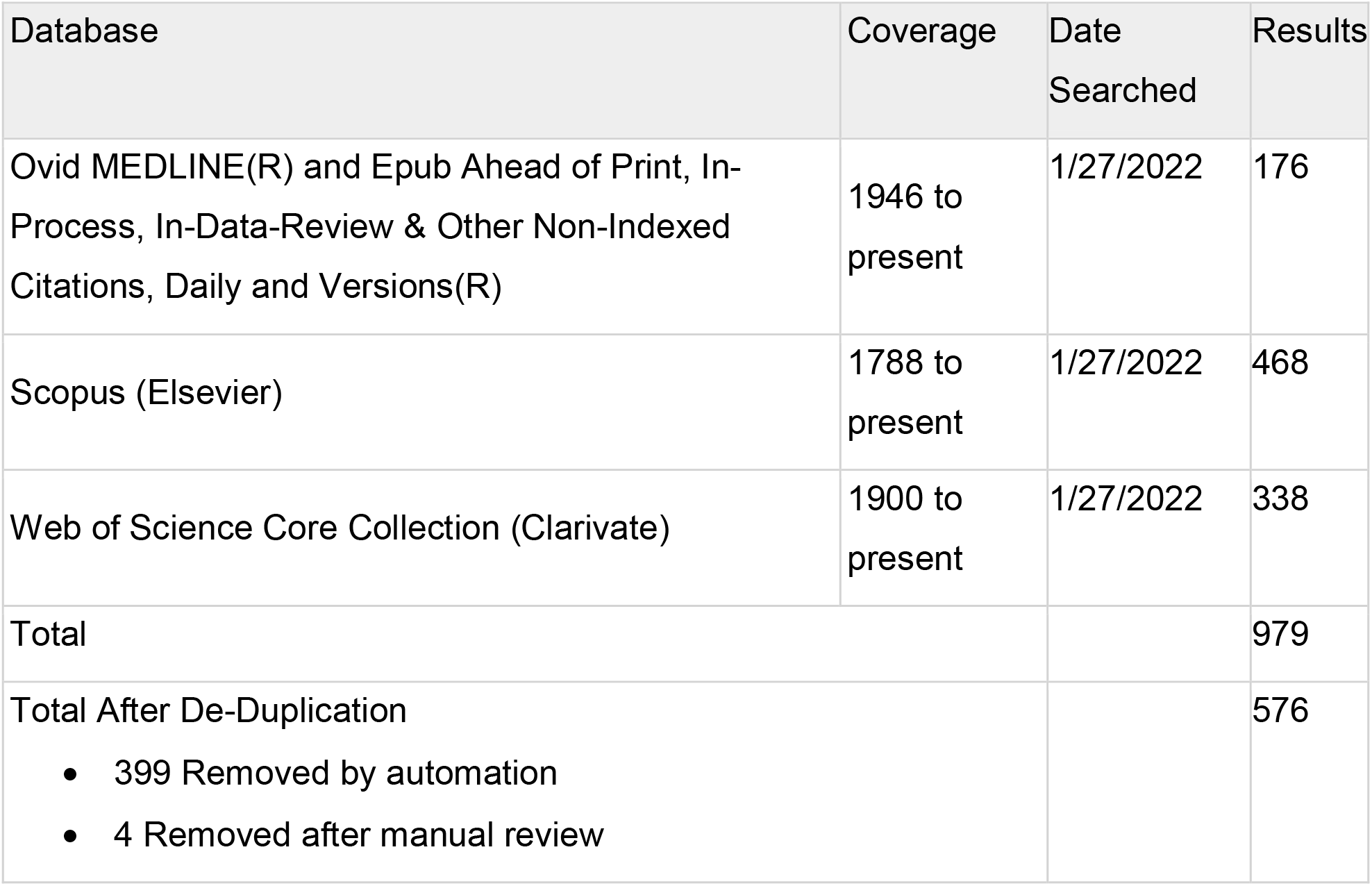

We searched the databases listed above on January 27, 2022. Records from each database were exported to EndNote. Duplicates were removed through the “Find Duplicates” function in EndNote and by manual review. Search strategies from each of the bibliographic databases are available below.

### Google Scholar

**Note:** Click on the hyperlinks below to see results from the Google Scholar searches Search 1 - <u>allintitle: CROT OR CROTs OR Datacite OR “CrediT taxonomy” OR “Contributor Role Ontology” OR “Contributor Role Taxonomy” OR TaDiRAH OR OpenVIVO</u>

Search 2 - <u>allintitle: authorship OR contributorship OR contributor AND credit OR credits OR contribution OR contributions</u>

Search 3 - <u>allintitle: author OR authors AND contribution OR contributions AND credit OR role OR roles</u>

Database(s): **Ovid MEDLINE^®^ and Epub Ahead of Print, In-Process, In-Data-Review & Other Non-Indexed Citations, Daily and Versions^®^** 1946 to January 26, 2022.

Search Strategy:

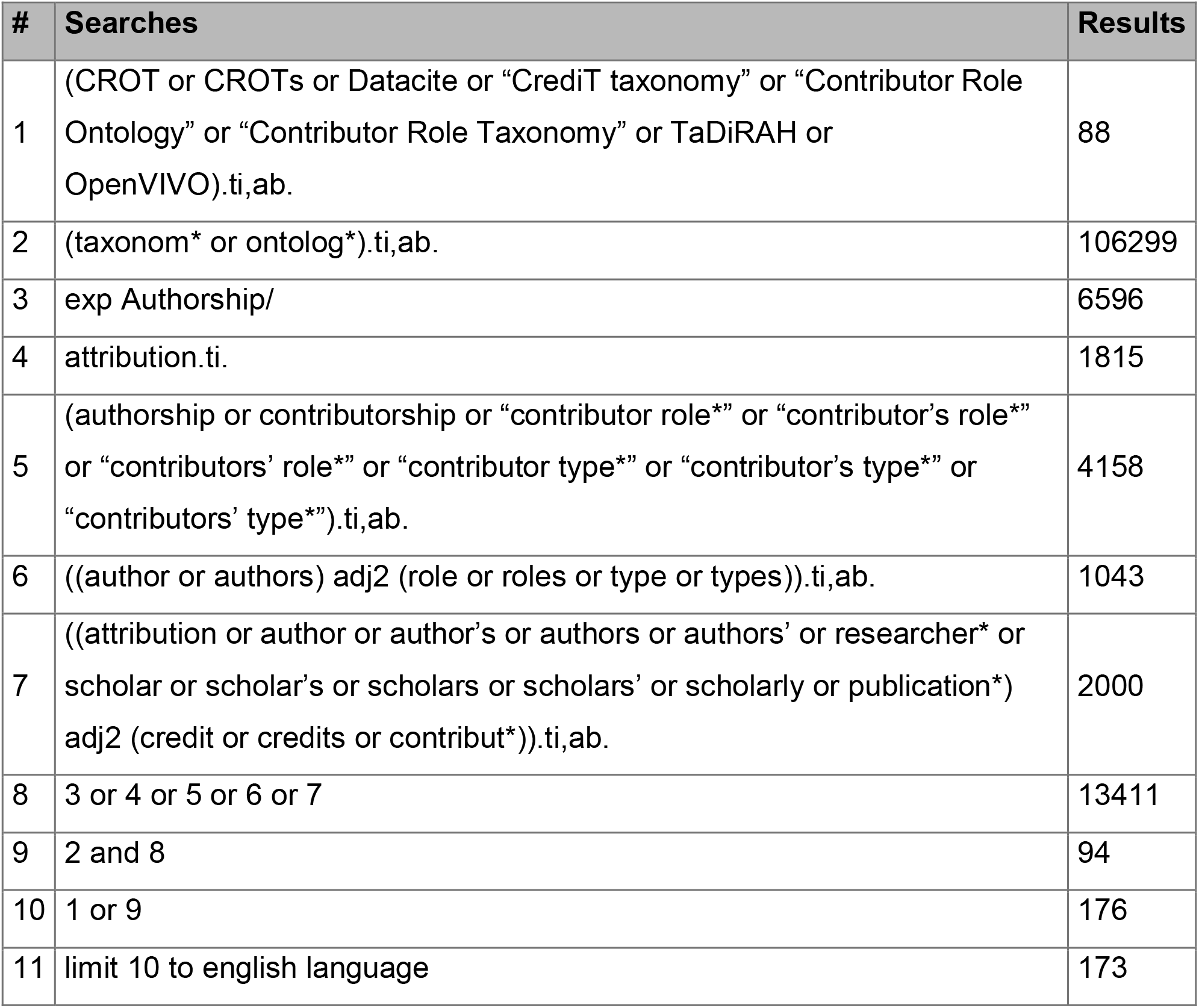

### Scopus

(TITLE-ABS-KEY(CROT or CROTs or Datacite or “CRediT taxonomy” or “Contributor Role Ontology” or “Contributor Role Taxonomy” or TaDiRAH or OpenVIVO)) OR ((TITLE-ABS(taxonom* or ontolog*)) AND ((TITLE(attribution)) OR (TITLE-ABS(authorship or contributorship or “contributor role*” or “contributor’s role*” or “contributors’ role*” or “contributor type*” or “contributor’s type*” or “contributors’ type*”)) OR (TITLE-ABS(((author or authors) W/2 (role or roles or type or types)))))) AND (LIMIT-TO (LANGUAGE,”English”)) AND (LIMIT-TO (DOCTYPE,”ar”) OR LIMIT-TO (DOCTYPE,”re”))

Limited to English, limited document type to Article and Review

### Web of Science

(TS=(CROT or CROTs or Datacite or “CrediT taxonomy” or “Contributor Role Ontology” or “Contributor Role Taxonomy” or TaDiRAH or OpenVIVO)) OR ((TI=(taxonom* or ontolog*) OR AB=(taxonom* or ontolog*)) AND (TI=(attribution) OR TS=(authorship or contributorship or “contributor role*” or “contributor’s role*” or “contributors’ role*” or “contributor type*” or “contributor’s type*” or “contributors’ type*”) OR TS=((author or authors) NEAR2 (role or roles or type or types))) OR TS=(((attribution or author or author’s or authors or authors’ or researcher* or scholar or scholar’s or scholars or scholars’ or scholarly or publication*) NEAR2 (credit or credits or contribut*))))

Refined By: Document Types: Articles or Review Articles | Languages: English

